# Impact of fungal hyphae on growth and dispersal of obligate anaerobic bacteria in aerated habitats

**DOI:** 10.1101/2022.03.03.482870

**Authors:** Bi-Jing Xiong, Sabine Kleinsteuber, Heike Sträuber, Christian Dusny, Hauke Harms, Lukas Y. Wick

## Abstract

Anoxic microsites arising in fungal biofilms may foster the presence of obligate anaerobes even in well-areated environments. Here, we analyzed whether and to which degree fractal hyphae of *Coprinopsis cinerea* thriving in oxic habitats enable the germination, growth, and dispersal of obligate anaerobic soil bacterium *Clostridium acetobutylicum.* Time-resolved optical oxygen mapping, microscopy and metabolite analysis revealed the formation and persistence of anoxic circum hyphal niches allowing for spore germination, growth and fermentative activity of the obligate anaerobe in an otherwise oxic environment. Hypoxic liquid films containing 80 ± 10% of atmospheric oxygen saturation around single air-exposed hyphae thereby allowed for efficient clostridial dispersal amid spatially separated (>0.5 cm) anoxic sites. Our results suggest that fungal biomass typical in soil (<550 μg g^-1^_soil_) may create anoxic microniches and enable activity as well as dispersal of obligate anaerobes near hyphae in an otherwise inhabitable environment.

## Introduction

Anoxic and hypoxic microsites foster the presence and activity of even obligate anaerobes in oxic environments (1–4) such as upland soils. As oxygen limitations typically arise if microbial oxygen consumption exceeds diffusive oxygen supply (1, 5), hypoxic microsites often coincide with hotspots of microbial activity such as the rhizosphere (6), the detritusphere (7), or biocrusts (8). Microbial biofilms (9–13) thereby often form steep oxygen gradients with oxygen depletion arising as shallow as ~80 μm beneath the air interface in mycelial layers (14, 15) as detected by needle-type oxygen microsensors (tip size: ~10 μm). Self-induced and spatially confined hypoxic microenvironments also form within filamentous fungal biofilms despite abundant spaces between hyphae and they are thought to contribute to fungal resistance to antifungal treatments (16). Anoxic fungal niches have also been observed to induce the growth of strict anaerobes (9, 10) suggesting that oxygen depletion may be a specific fungal mechanism to modulate the mycosphere chemistry (10, 17). Typically forming 0.05-1 mg of biomass dry weight (biomass, dw) per g soil, fungi (18–20) may embody up to 75% of the subsurface microbial biomass (19). Being predominantly aerobic (21) they consume oxygen at rates of up to 180 nM_oxygen_ min^-1^ mg_biomass,dw_^-1^ (14). Unlike in tightly packed industrial biofilms, soil fungi often develop extensively fractal mycelia that allow them to access heterogeneously distributed nutrients and carbon sources (22) and to bridge mycelial source- and sink-regions (23). Forming hyphae with lengths of ≈10^2^ m g^-1^ in arable and up to 10^4^ m g^-1^ in forest topsoil (19), mycelia thereby also serve as important pathways for bacterial dispersal (‘fungal highways’) (24) enabling the colonization of new habitats (25–28), horizontal gene transfer (29), or predation (30). Expressing hydrophobic cell-wall proteins (hydrophobins), hyphae thereby overcome air-water interfaces and bridge air-filled pores with nutrient-rich aqueous zones containing little or no oxygen. In upland soils furthermore, conditions may switch rapidly between oxia and anoxia (1) (e.g. in response to heavy rainfall or waterlogging), leading to a rapidly changing distribution of oxygen. As fungi may both form and bridge anoxic microsites, we here assess to which degree hyphae thriving in oxic habitats enable spore germination, vegetative growth, and dispersal of strictly anaerobic bacteria. Towards this aim, we determined spatial and temporal oxygen profiles around the filamentous mycelia of the fast-growing (~100 μm per h (31)) aerobic soil fungus *Coprinopsis cinerea.* Planar optodes and custom-made micrometer-sized oxygen-sensitive beads determined mycelial oxygen distribution. The sporeforming soil bacterium *Clostridium acetobutylicum* was chosen as a representative of strict anaerobes due to its inability to grow, ferment and swim under oxic conditions (32). Its oxygen-tolerant spores only germinate into fully active vegetative cells under anoxic conditions (33). Spatially and temporally resolved oxygen mapping revealed the formation and persistence of anoxic regions in air-exposed *C. cinerea* mycelia that enabled spore germination and growth of *C. acetobutylicum.* For the first time, we also document active longdistance (>0.5 cm) dispersal of an obligate anaerobe along airexposed hyphae between two anoxic sites. Our results demonstrate that the occurrence and activity of aerobic mycelia can create anoxic microniches and facilitate activity and spatial distribution of obligate anaerobes in an otherwise oxic environment. This enlarges our understanding of the ecological role of the mycosphere (i.e. the microhabitat surrounding and affected by hyphae and mycelia) (34) for microbial dynamics and functional stability at oxic-anoxic interfaces in heterogeneous ecosystems.

## Materials and Methods

### Microorganisms, medium and growth conditions

The obligate anaerobe *C. acetobutylicum* DSM 792 (type strain, purchased from DSMZ – German Collection of Microorganisms and Cell Cultures, Braunschweig, Germany) was cultivated for three days at 37°C under strictly anoxic conditions with an N_2_ headspace in 200mL serum bottles containing 50 mL of SM824 medium (35). Using a syringe, two mL of the culture with an optical density OD_600_ = 0.81 were then removed, centrifuged at 4,000 × g at 10°C for 5 mins, the supernatant discarded, the cells re-suspended in 2 mL air-saturated SM824 medium and then used as inoculum (cf. below). The remaining culture was further cultivated until endospore formation was observed (typically at t = 7 d, Fig. S1, SI). *C. acetobutylicum* spores were obtained as described by Yang et al. (36) (cf. SI) to form inocula of an OD_600_ <0.02 in SM824 medium (cf. below). The basidiomycete *C. cinerea* strain AmutBmut (37) served as filamentous fungus (38). It was cultivated at 30■°C for three days on yeast-malt extract-glucose medium (39). Using a scalpel, a small piece (Ø <1 mm) of *C. cinerea* inoculum was cut from the peripheral growth zone and used as inoculum in the microcosms.

### *Time-lapse*, in vivo *mapping of mycelial oxygen profiles*

#### Microcosm

All microcosms were handled and used under laboratory atmosphere conditions. Microcosms for mycelial oxygen profile mapping consisted of a fungus-inoculated agarose pad (Ø 18 mm, h: 1 mm; Fig. 1a) that was inversely placed on an oxygen optode (Ø 18 mm, SF-RPSu4, PreSens, Regensburg, Germany; Fig. 1b) as described earlier (31). Briefly, 400 μL of aerated SM824 medium (1.5% low-melt agarose, Karl Roth, Karlsruhe, Germany) were placed on a circular cover slide (Ø 18 mm, ibidi, Gräfelfing, Germany), immediately covered by a second cover slide, and allowed to cool for 10 min. After removing the top slide with tweezers, the agarose pad was centrally inoculated with a small piece (Ø <1 mm) of fungal inoculum, and incubated for 48 h at 30°C. The pad was then flipped over and attached to an oxygen optode that itself was glued to the glass bottom of a Petri dish (*μ*-Dish 35 mm, low, ibidi). The second cover slide was removed immediately. Five sterile agarose pads (Ø 8 mm) cut from a 1-cm thick SM824 agar plate were evenly placed around the inoculated pad to keep the microcosm moisturized.

**Figure 1.**
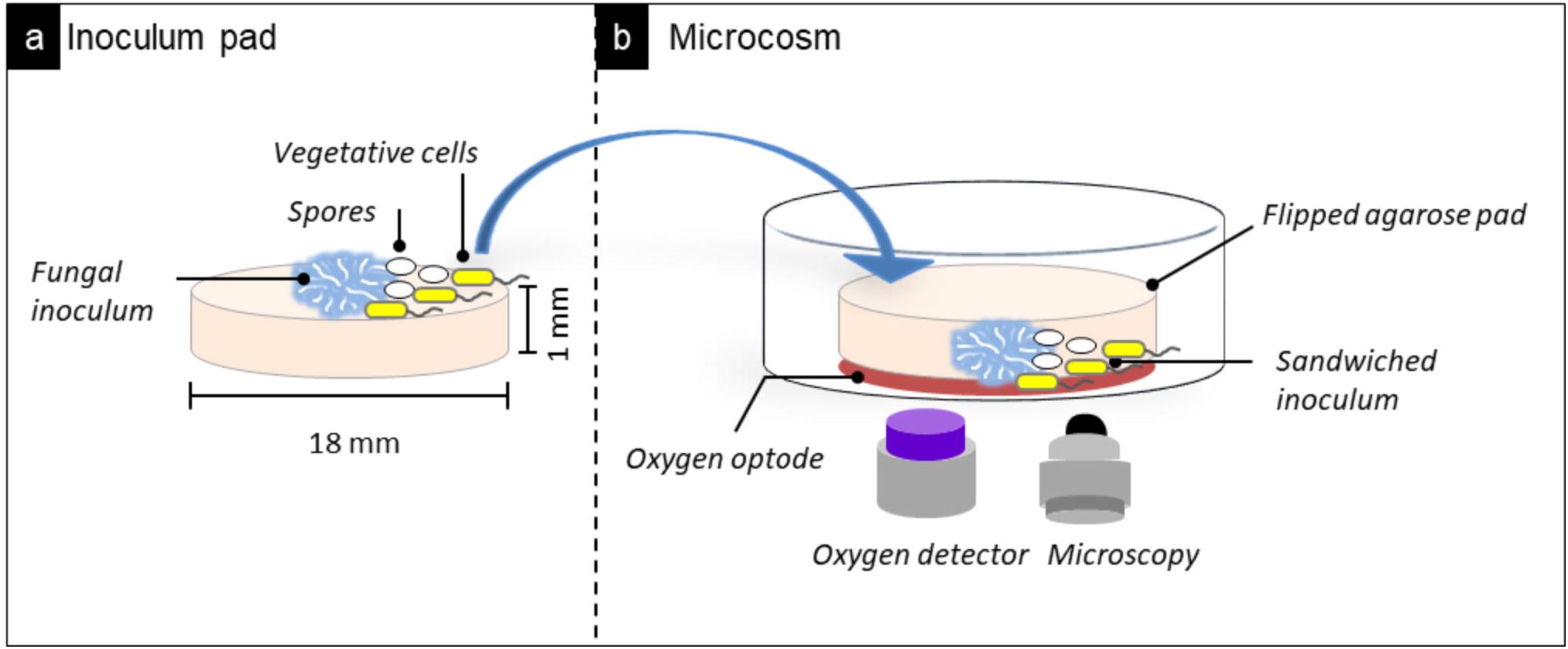
Schematic view of the microcosm used for *in vivo* time-lapse mapping of mycelial oxygen profiles (experiment 1) and the detection of mycelia-induced spore germination and bacterial growth (experiment 2). **a)** An agarose inoculum pad that was inverted and placed in the **b)** microcosm to sandwich the fungal inoculum between an oxygen optode at the bottom and the overlying agarose pad. The oxygen optode is glued to the glass bottom of a Petri dish and sensor signals are monitored by a commercial detector. The development of the fungal networks on the optode surface is imaged by bright-field microscopy. In experiment 2, vegetative cells or spores are placed at the center of agarose inoculum pad or 3 mm away from the fungal inoculum respectively. Growth, spore germination and activity of obligate anaerobe are followed by microscopy.

#### Mycelial growth and oxygen distribution in the mycosphere

Immediately after placing the pad (t = 0 h), oxygen profiles around the fungal inoculum (area: 1.8×2.5 mm) were monitored for 15 min at intervals of 30 s, using a commercial detector unit (VisiSens TD Detector Unit DU02, PreSens). Combining microscopic observation (Nikon AZ100, Amsterdam, The Netherlands) and optical sensing techniques the mycelial development and oxygen concentrations in the whole pad were then mapped at t = 0.3, 48, 72, 120 and 200 h. As the commercial oxygen sensing unit did not support cm-scale sensing, the microcosm was fixed to a connection arm (Edelkrone, Langen, Germany) that was mounted on a software-controlled microscope stage (NIS-ELEMENTS, Basic Research, Nikon). To map the agarose pad with an area of ca. 250 mm^2^, >100 spots of 1.8×2.5 mm were analyzed and stitched using open-source software ImageJ (https://imagej.nih.gov/ij/). An interval of 3000 ms was allowed between shifting two spots. In this period, the oxygen profile at a given spot was measured with the detector unit. After oxygen mapping (duration: ~15 min), bright-field microscopic images of the mycelium were taken at each time point using the large image acquisition function of the microscope (15 × magnification). The microcosms (Fig. 1b) were stored at 30°C in the dark between sampling times. Triplicate experiments were performed. The calibration (Fig. S2) of the oxygen optode is described in the SI.

### *Hypha-induced vegetative growth and spore germination of* C. acetobutylicum

Similar microcosms as described above (Fig. 1b) yet without the semi-transparent oxygen sensor served to examine the vegetative growth and fermentative activity of *C. acetobutylicum* in presence of *C. cinerea:* 1 μL of a *C. acetobutylicum* suspension (OD_600_ = 0.81, or 8.1×10^7^ cells mL^-1^) was inoculated to the centre of the agar pad that had been pre-incubated with *C. cinerea* for 48 h. Then the pad was flipped over and attached to the glass bottom of a Petri dish (*μ*-Dish 35 mm, low, ibidi). To moisturize the air in the Petri dish, five sterile agarose pads (Ø 8 mm, h: 1 cm) were evenly placed around the inoculated pad. The microcosms were incubated as described above and examined daily with a microscope. At t = 7 d the microcosms were harvested for i) DNA extraction and subsequent 16S rRNA gene sequencing, ii) total and viable bacterial cell counting, and iii) analysis of water-soluble metabolites (cf. SI). Sterile pads and pads inoculated with *C. cinerea* only or *C. acetobutylicum* only served as controls. All experiments were performed in triplicate. DNA was extracted using a commercial extraction kit (DNeasy PowerSoil KitKit, Qiagen, Hilden, Germany) following the manufacturer’s protocol, and the 16S rRNA gene was partially sequenced using the primers of 27f and 519r as described elsewhere (40). For total and viable cell counting, bacterial cells in the microcosms were recovered as described in the SI. Total and viable bacterial cell counting was done using an automated microbial cell counter (QUANTOM Tx^TM^, Logos, Gyeonggi, South Korea) following manufacturer’s protocol.

For spore germination experiments, 3 μL of spore suspension (OD_600_ <0.02) were inoculated to the agarose pad (Fig. 1a) at a distance of ~3 mm from the fungal inoculum and the microcosm was incubated in the dark for 24 h at 30°C under laboratory atmosphere conditions. After 24 h, spore germination and the subsequent cellular growth were monitored by phase contrast microscopy (exposure time 300 ms, LED light source intensity 4.7 V) at 20 mins intervals for 36 h. During the microscopic imaging, the microcosm was incubated at constant temperature (30°C) by using a heating system (XLmulti S2, Carl Zeiss Microscopy GmbH, Jena, Germany) mounted to the microscope (Axio Observer, Carl Zeiss Microscopy GmbH). At the end of the experiments, the microcosms were harvested for DNA extraction and 16S rRNA gene sequencing as described above.

### *Dispersal of* C. acetobutylicum *along hyphae*

Two 1-mm thick agarose pads were inoculated with *C. cinerea* (cf. above, Fig. 1a) and incubated at 30°C for 48 h. After that, 1 μL of vegetative *C. acetobutylicum* cells (OD_600_ = 0.81) was placed onto the fungal inoculum in one agarose pad. Immediately thereafter, the fungus-bacteria-inoculated pad (agar pad A) and exclusively fungus-inoculated pad (agar pad B) were both flipped over and placed at a distance of 5 mm on a microscopy slide (26×76 mm, ibidi). The microscopy slide was then transferred to a plastic Petri dish (90 mm, Thermo Fisher, Waltham, USA), and five circular agarose pads (Ø 10 mm, h: 1 cm; SM824 agar) were evenly placed around the slide to keep the microcosm moisturized during incubation for 7 d at 30°C and ambient air. All microcosms were prepared in triplicate and monitored daily by microscopy to examine the formation of hyphae between two agar pads and the presence of *C. acetobutylicum* cells moving along them, respectively. To exclude potential bacterial contamination, DNA was extracted from both agar pads at t = 7 d and sequenced as described above.

### Hyphal oxygen mapping by lifetime-based oxygen-sensitive beads

Oxygen content in the liquid film surrounding *C. cinerea* hyphae was also measured using custom-made, lifetime-based oxygen-sensitive beads (Ø = 8 μm). *C. cinerea* was inoculated to the 1-mm thick agarose pad and incubated at 30°C for 48 h. The oxygen-sensitive beads were dispersed in sterile deionized water and three drops (1 μL each drop) suspension were placed at ~2 mm distance from growing hyphal tips to the agar surface. The agarose pad was then incubated overnight at 30°C to let the hyphae overgrow the beads. Position and lifetime-based luminescence quenching of the ruthenium-phenanthroline-based phosphorescence dye in individual oxygen beads were measured using an OPAL system (Colibri Photonics, ibidi) connected to an automated inverted Zeiss microscope (Axio Observer, Carl Zeiss Microscopy GmbH). The 532 nm LED of the OPAL and a filter cube with 531/40 nm (excitation), 607/70 nm (emission) were inserted in the optical light path during measurements. The SI describes the calibration of the oxygen beads.

Swimming activity of *C. acetobutylicum* in liquid SM824 medium with different oxygen levels was examined microscopically as described in SI.

## Results

### *Time-resolved* in vivo *mycelial oxygen distribution*

Combining microscopic observation and optical oxygen sensing, we mapped growth and spatio-temporal oxygen concentrations in aerated mycelia of *C. cinerea* (Fig. 2, Video S1). Zones of oxygen depletion formed at the inoculation point as shortly as <6 min (Video S1) after attaching the pad to the oxygen optode (Fig. 1b). Diameters of oxygen-free zones then steadily expanded from 2.3 ± 0.8 mm at t = 0.3 h to 18 ± 0.0 mm at t = 120 h before decreasing to 2.3 ± 0.6 mm at t = 200 h (Fig. S3, SI). At the mycelial borders we observed steep oxygen gradients from 100% to 0% air saturation over a distance of ~2 mm (Fig. 2, row c).

**Figure 2.**
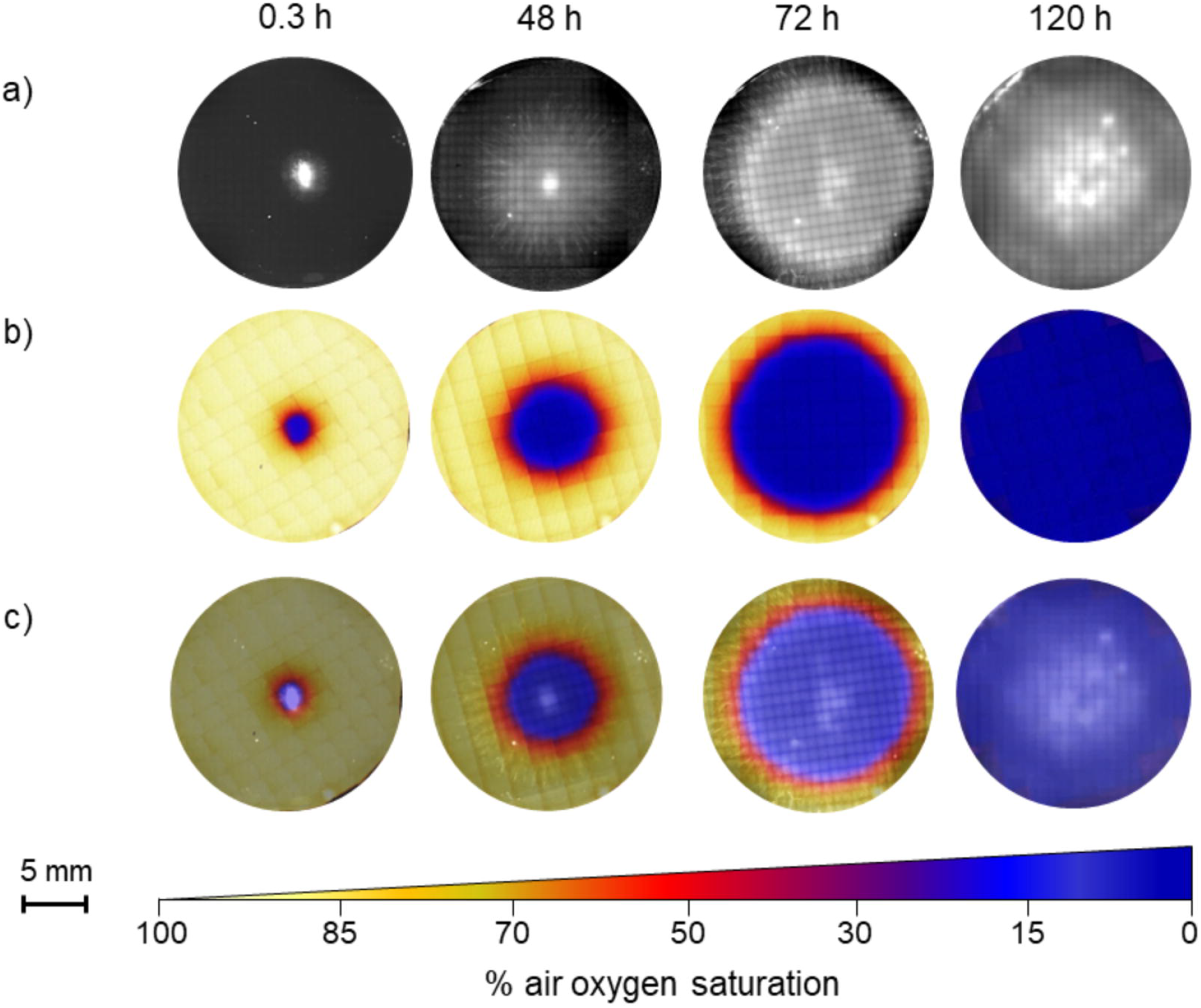
Growth and *in vivo* time-lapse mycelial oxygen profiles of *C. cinerea*. (cf. Video S1). Rows **a** & **b** depict micrographs of the development of mycelial growth and mycelial oxygen profiles of *C. cinerea* monitored over 120 h at 30°C using an oxygen optode. Row **c** shows an overlay of optode and bright-field micrographs. Darkblue and light-yellow colors refer to ca. 0 – 100% of air oxygen saturation in water at 30°C. (cf. color bar).

### *Growth and fermentative activity of* C. acetobutylicum *in presence of mycelia*

To validate the presence of the optically mapped oxygen-free zones, we inoculated *C. acetobutylicum* cells to *C. cinerea* and tested if mycelial activity enables growth of the obligate anaerobe. After 7 d of air-exposed growth in presence of mycelia, *C. acetobutylicum* formed dense colonies in the mycosphere (Fig. S4a, t = 7 d) accounting for a >1300-fold increase in cells numbers from 3.5×10^5^ ± 3.3×10^3^ (t = 0 d) to 4.8×10^8^ ± 3.5×10^7^ cells g^-1^_agat_ = 7 d (Fig. S4c). Micrographs revealed typical traits of *C. acetobutylicum* including rod-shaped cells and the formation of optically brighter regions at one pole (Fig. S4a, t = 7 d) as signs of endospore formation. Comparison of total and viable cell counts (Fig. S4c) further showed that >75% of the total cells were viable at t = 7 d. DNA extraction and subsequent 16S rRNA gene sequencing confirmed that *C. acetobutylicum* was the only bacterium present in the microcosms as the partial sequence was identical to the published 16S rRNA gene of *C. acetobutylicum* (acc. no. NR_074511). In the absence of mycelia, no growth of the obligate anaerobes was observed (Fig. S4b) with cell numbers that were too low to be detected by the cell counter (Fig. S4c). Bacterial growth in presence of mycelia went along with the production of significant amounts of butyrate and 1-butanol, i.e. typical water-soluble products of clostridial fermentative metabolic activity (Fig. S5, SI). No such metabolites were found in controls with only *C. acetobutylicum* or *C. cinerea* inocula under aerobic conditions.

### *Hypha-induced germination of* C. acetobutylicum *spores*

Microscopic observation of *C. acetobutylicum* germination near active growing *C. cinerea* hyphae further evidenced the presence of anoxic niches in the mycosphere (Fig. 3, Video S2). Time-lapse microscopic imaging revealed a phase-bright appearance of dormant spores (Fig. S1, SI) in the absence of hyphae at t = 0-30 h (Video S2). At t = 30-32 h (Fig. S6) the halo surrounding of the spores gradually increased and spores lost their phase-bright appearance between t = 31 h 40 min and t = 32 h, i.e. when the first hyphal tip became visible at a distance of 13 μm from the spore (Fig. 3). At t = 32 h 20 min, the shedding of the spore finished and the formation of a dense bacterial mat became visible (t = 32 h 20 min – 60 h, Fig. 3). The mat was demonstrated to consist of *C. acetobutylicum* by 16S rRNA gene sequencing. In the absence of hyphae, no clostridial spores germinated (Fig. S7).

**Figure 3.**
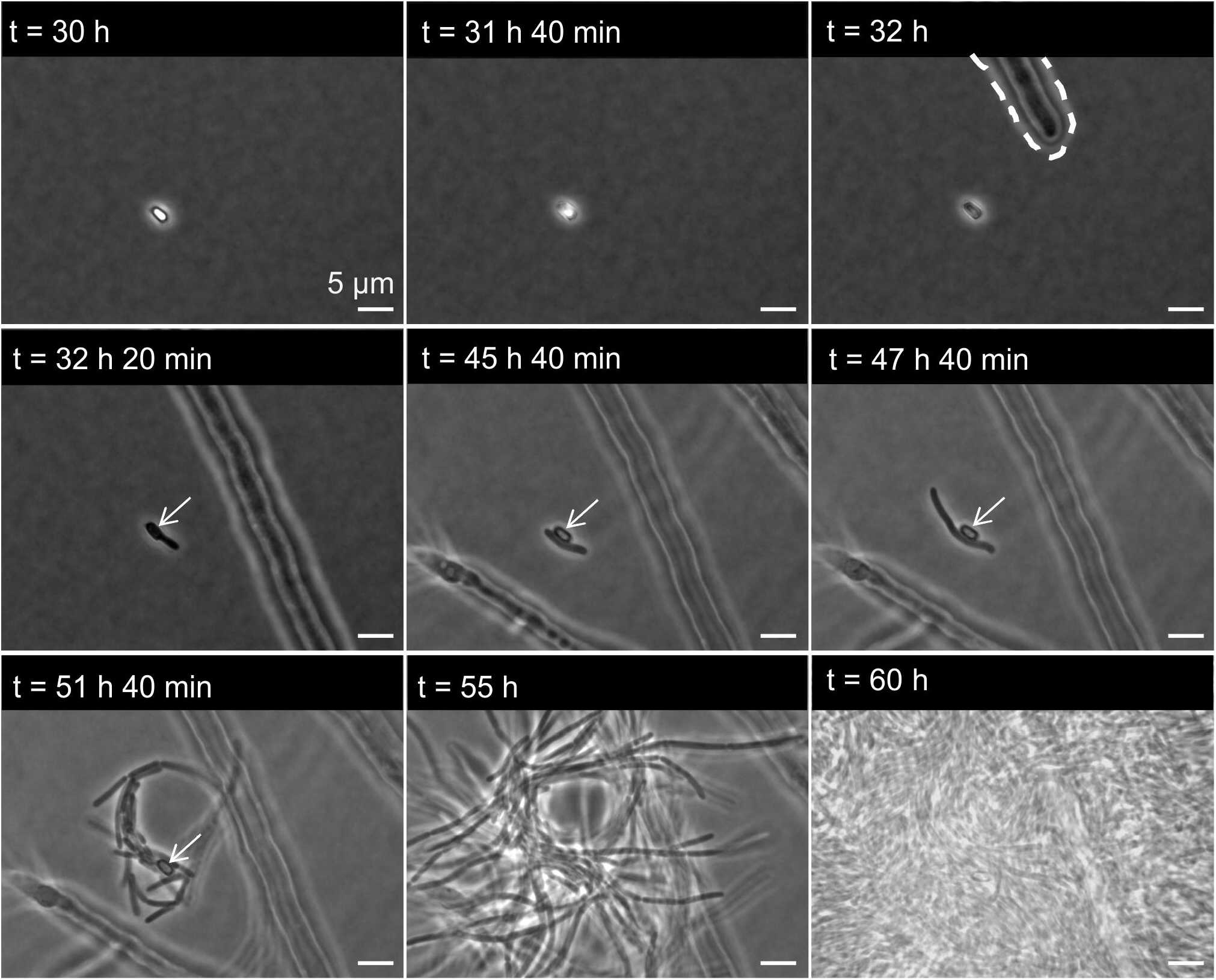
Germination and growth of *C. acetobutylicum* spores in the hyphosphere of *C. cinerea*. At *t* = 30 h: phase-bright appearance of a dormant spore initially inoculated ca. 3 mm from the hyphal tip. *t* = 31 h 40 min: swelling of spore; one pole of the spore is less bright. *t* = 32 h: appearance of hyphae and spore without phase-bright appearance. *t* = 32 h 20 min: appearance of germ tube. *t* = 45 h 40 min: longitudinal growth phase is completed. *t* = 45 h 40 min – 60 h: symmetric cell division and formation of bacterial colony. White arrows point at empty spore shells after germination.

### *Dispersal of* C. acetobutylicum *along air-exposed mycelia*

*C. acetobutylicum* cells growing in the mycosphere of *C. cinerea* in the agar pad A (Fig. 4a) remained motile (Video S3) and started to colonize and disperse along hyphae linking the two nutrient-rich agarose pads (Fig. 4a) at t = 5 d (Fig. 4b, Video S4). Such dispersal resulted in the colonization of agar pad B (Fig. 4a) and subsequent growth of *C. acetobutylicum* therein (Fig. S8). In the absence of hyphae, no clostridia between the two agar pads and no cells in agar pad B were observed. In a separate experiment, we further analyzed the effect of oxygen on swimming of *C. acetobutylicum.* Micrometer-sized lifetime-based oxygen-sensitive beads near air-exposed hyphae revealed that oxygen levels decreased to 73 ± 5% air saturation (Fig. 4d, 0-1 μm) at the direct hyphal surface, 85 ± 3% in the apparent liquid film at distances of 1-10 μm above the hyphal surface (Fig. 4d, 1-10 μm), and 100 ± 1% outside the hyphal liquid film (Fig. 4d). To study the effect of oxygen on clostridial motility we microscopically assessed dispersal of *C. acetobutylicum* cells in liquid medium with 80% and 100% air-saturated oxygen concentrations: 2.2%, 1.2% and 0.8% of the observed cells actively swam after 1, 15, and 30 min in ~80% air saturated media (Table S1). No active swimming was observed, however, in a fully air-saturated environment (Table S1).

**Figure 4.**
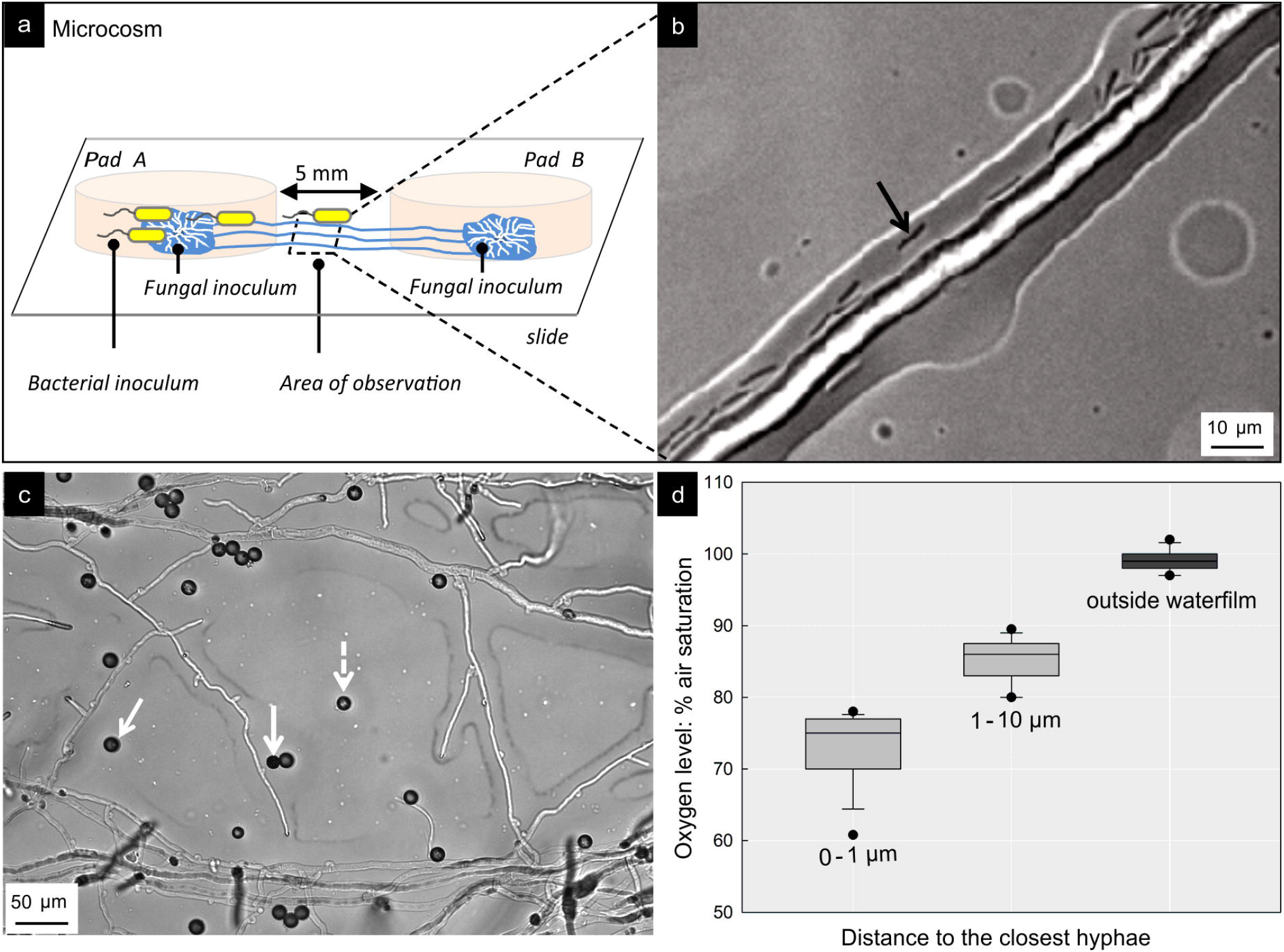
Spatial dispersal of *C. acetobutylicum* along *C. cinerea* hyphae. **a**, Schematic view of the microcosm used for time-lapse monitoring of dispersal of *C. acetobutylicum* cells along air-exposed hyphae of *C. cinerea.* Agar pad A was inoculated with *C. cinerea* and *C. acetobutylicum* and pad B with *C. cinerea* only. **b**, Micrograph of *C. acetobutylicum* in air-exposed hyphae of *C. cinerea* between agar pads A & B at t = 5 d. **c**, Micrograph of representative distribution of oxygen-sensitive beads (Ø 8 μm) between hyphae of *C. cinerea* overgrowing an agar surface. White arrows exemplify the presence of beads either immersed in the hyphal liquid film extending to ca. 10 μm around hyphae. **d**, Distribution of sensed oxygen concentration by oxygen-sensitive beads in apparent distance from hyphal surface (80 ± 7.4% air oxygen saturation, 0-10 μm, *n* = 53; 99 ± 1.5% air oxygen saturation, >10 μm, *n* = 13).

## Discussion

### Mycelia form anoxic niches and enable activity of anaerobes

By combining oxygen sensing and culture-based approaches, our study shows that growing mycelia of *C. cinerea* form anoxic niches (Fig. 2) allowing for development (Figs. 3 & S4) and metabolic activity (Fig. S5) of the obligate anaerobic bacterium *C. acetobutylicum.* Although few studies report anoxic niches in fungal biofilms (10, 16, 17), effects of fungal oxygen consumption on bacterial-fungal interactions are still poorly described. Biofilms of oxygen consumption by the yeasts candida created anaerobic niches in the oral cavity (17), thereby influencing the composition of its associated microbiome by favoring anaerobic bacteria over aerobes (26, 41). Being prevalent in heterogeneous terrestrial habitats fungi often form dense mycelial mats on dead organic material or exist in fungal-bacterial biofilms. Metabolically active fungi are thereby likely to create spatially distinct anoxic/hypoxic niches. Assuming typical fungal oxygen consumption of 1.63×10^-3^ mol m^-3^ s^-1^ (15, 42, 43), a rough calculation (for details cf. SI) indicates that ~543 μg g^-1^_soil_ of fungal biomass would suffice to create anoxia if the mycelia were covered by a water film of 1 mm thickness. Given a typical fungal biomass density of 50-1000 μg g^-1^_soil_ (18–20), our calculation suggests that aerobic fungi may play an overlooked role in creating (and bridging; cf. below) anoxic micro-niches in otherwise oxic habitats. Such assumption is in line with the observed formation of anoxic habitats at the periphery of oxygen consuming fungal-bacterial hotspots in soil (5, 6, 44). It also supports the previously described co-occurrence of aerobes and anaerobes within microbial communities in aerated soils (4, 45) or the presence of genes responsible for the anaerobic biosynthetic pathways in the mycosphere (7, 8).

### Mycelia mediate activity and spatial organization of anaerobes in air-exposed habitats

While the mycosphere is known as typical habitat for aerobic bacteria (46, 47), our study reveals that hyphae may also evoke the germination, activity and growth of spores of anaerobic *C. acetobutylicum* in an aerated habitat (Fig. 3, Video S2). Spore germination of *C. acetobutylicum* thereby started already at a distance of ~13 μm from the hyphal tip (Fig. 3). This suggests that hyphal oxygen consumption affected similar regions as described for pH changes (31), enzymatic activity (48) or the dispersal of bacteria (29, 49). Air-exposed hyphae of *C. cinerea* by this means also enabled the dispersal of *C. acetobutylicum* and the colonization of anoxic habitats separated by ambient air (Fig. 4, Video S4).

Even though fermentation and growth of *C. acetobutylicum* is immediately halted by the presence of oxygen (32), literature shows that *C. acetobutylicum* (50) and other strict anaerobes adapt to oxygen exposure by using defense mechanisms to minimize oxygen stress (2, 51). Our experiments (Table S1) for instance revealed that ca. 1% of *C. acetobutylicum* cells remained up to 30 min motile if exposed to 80% air-saturated medium (Table S1); i.e. oxygen levels similar to those observed by oxygen-sensing microbeads in a liquid film formed around hyphae (Fig. 4d). As clostridia characteristically express highest motility in the exponential growth phase (52), the most active cells may thus move along hypoxic hyphae. Assuming a mean swimming speed of *v* = 5-25 μm s^-1^ (53, 54) and active swimming for t = ~30 min, hyphosphere clostridia would be able to disperse over distances *s* = *v* × *t* of 27-45 mm between anoxic niches.

Our observation that obligate anaerobe *C. acetobutylicum* actively swim in the hypoxic liquid around hyphae suggests that mycelia may not only mediate the transfer of aerobes (24) but also anaerobes (Fig. 4) between spatially separated habitats (as summarized in Fig. 5). Such finding seems particularly important for soil, as soil microbial communities are constantly exposed to fluctuating conditions (55, 56) that can lead to physico-chemically distinct zones, habitat patchiness, and the formation of hotspots of microbial activity. Localized microbial activity likewise may lead to the formation of oxygen gradients (5) and spatial separation of aerobes and anaerobes (57). Fractally growing fungi in soil (22) thereby may often overgrow and link habitats with differing oxygen concentrations. Due to their ability to cope with short-term hypoxic conditions (e.g. by the use of fermentative metabolism (58), fungal networks may thus stably bridge oxic, oxic-anoxic, or anoxic interfaces (Fig. 5) even under varying environmental conditions. Given the short-term tolerance of strict anaerobes to oxygen and presumed reduced oxygen contents in the hyphosphere, we propose that hyphae in spatially heterogeneous environments may serve as a good network for the activity and spatial organization of a wide range of bacteria: including obligate anaerobes, facultative anaerobes, aero-tolerant anaerobes, micro-aerophiles and aerobes. Exploring mycosphere processes beyond traditionally assumed boundaries will further unravel the interactions, functioning and stability of fungal-bacterial communities in microbial ecosystems.

**Figure 5.**
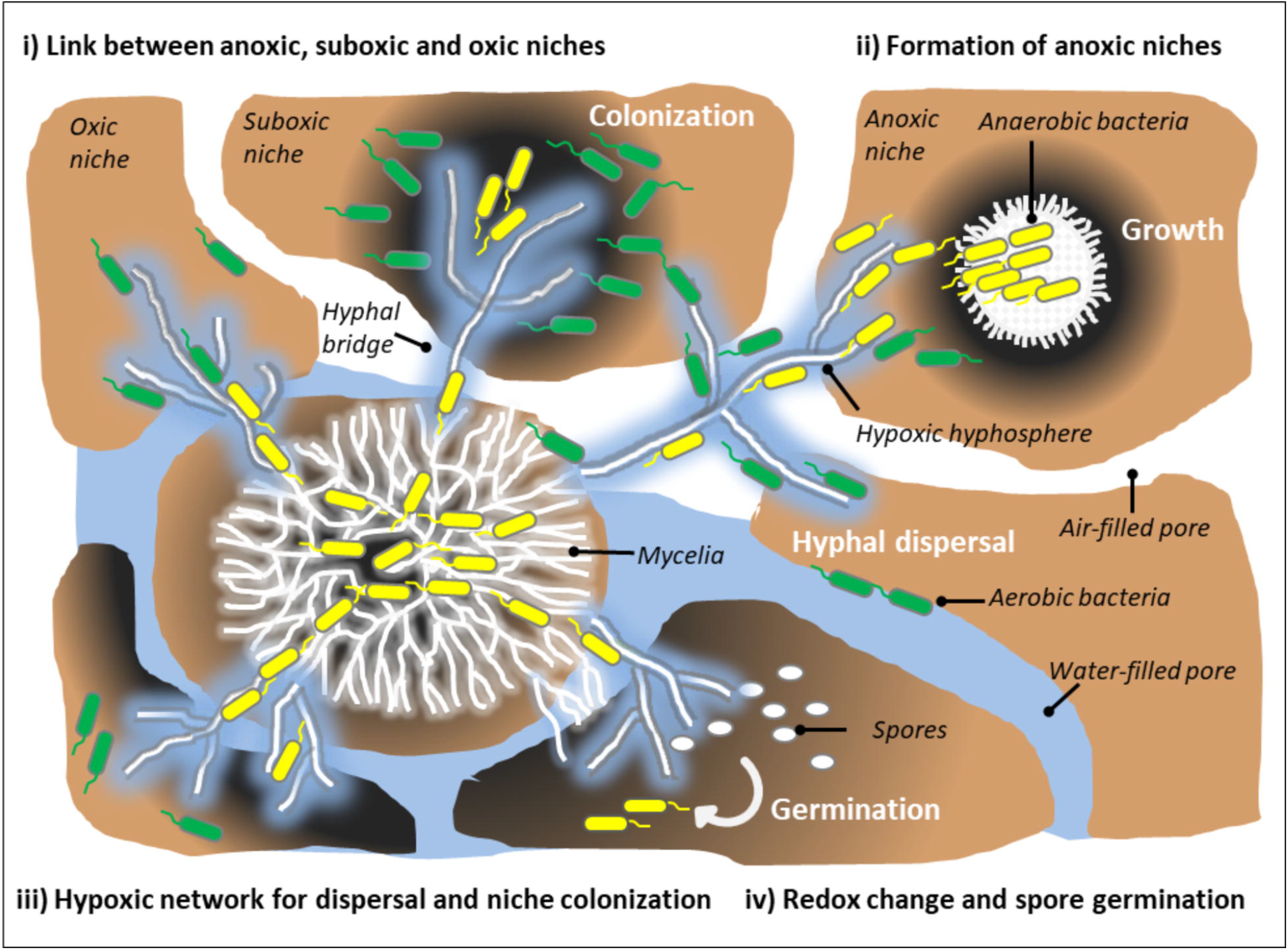
Mycelia mediate activity and spatial organization of anaerobic microorganisms. By forming extended fractal networks, filamentous fungi are well-adapted to heterogeneous soil environments, which often may form microbial hotspots and anoxic/hypoxic microniches. Due to their fractal structure and ability to cope with short-term hypoxic conditions, fungi may form stable networks that form hyphal links between anoxic/hypoxic, oxic and/or oxic-anoxic niches in soil (i). Being predominantly aerobic, fungal mycelia may consume oxygen at higher rates than can be delivered by their environment. This leads to the formation of mycelial anoxic niches that enable growth and activity of anaerobic bacteria (ii). Active hyphae also reduce the oxygen content in their hyphosphere. Even if air-exposed, mycelia hence may form a hypoxic network for dispersal of anaerobic and aerobic bacteria that enable the colonization of new niches (iii). Reduction of hyphosphere oxygen content and changes of concomitant redox conditions also stimulate the activity of anaerobes as reflected, for instance, by the germination of their spores (iv). Dark grey color in the figure refers to reduced oxygen level.

## Acknowledgement

The authors acknowledge financial support by the China Scholarship Council (CSC), the German Academic Exchange Service (DAAD) in the frame of the FungDeg project and the Collaborative Research Centre AquaDiva of the Friedrich Schiller University Jena, funded by the Deutsche Forschungsgemeinschaft (DFG, German Research Foundation)-SFB 1076-project number 218627073 and the Helmholtz Centre for Environmental Research – UFZ. The authors wish also to thank the professional technical support from Kristin Lindstaedt, Claudia Heber and Ute Lohse.

## Author contributions

BX and LYW designed the study. BX performed the experiments. BX, LYW, SK, HS, CD and HH wrote the manuscript. All authors read, revised and approved the final manuscript.

