## Supplemental Information for "Impact of fungal hyphae on growth and dispersal of obligate anaerobic bacteria in aerated habitats"

**SUPPLEMENTARY INFORMATION**

**Fungal hyphae enable spore germination, growth and dispersal of obligate anaerobe *Clostridium acetobutylicum* in oxic habitats**

Bi-Jing Xiong^1^,	Sabine Kleinsteuber^1^, Heike Sträuber^1^, Christian Dusny^2^, Hauke Harms^1^, and Lukas Y. Wick^1*^

^1^ Department of Environmental Microbiology, Helmholtz Centre for Environmental Research – UFZ, Permoserstraße 15, 04318 Leipzig, Germany.

^2^ Department of Solar Materials, Helmholtz Centre for Environmental Research – UFZ, Permoserstraße 15, 04318 Leipzig, Germany.

This supplement contains:

- Information on ‘Materials and Methods’
- 1 Table
- 8 Figures

^*^ Corresponding author: Mailing address: Helmholtz Centre for Environmental Research – UFZ. Department of Environmental Microbiology, Permoserstraße 15, 04318 Leipzig, Germany. phone: +49 341 235 1316,.

**Materials and Methods**

**Harvest and purification of spores**

To harvest *C. acetobutylicum* spores, 20 mL of the sporulated culture was centrifuged at 12,850 × g for 10 min at 4^o^C, the pellet was resuspended in 20 mL sterile deionized water and centrifuged again as described above. The pellet was then resuspended in phosphate-buffered saline (PBS buffer, pH 7.4) containing 500 µg mL^-1^ lysozyme and sonicated for 5 min to release the spores from the mother cells. The suspension was then incubated at 37^o^C for 2 h to digest the vegetative cells and centrifuged at 12,850 × g for 10 min at 4^o^C. The pellet was suspended with 20 mL deionized water and the vegetative cell debris was removed by ten times repeated washing and centrifugation (2,050 × g, 20 min) steps. Thereby, spore suspension with an OD_600_ = 0.41 was prepared. The spore suspension was 1000 × diluted with sterile water (final OD_600_ < 0.02) before inoculation to the microcosms.

**Calibration of the oxygen sensor optode**

A conventional two-point calibration in an oxygen-free and an air-saturated environment, respectively, was performed to calibrate the oxygen sensing optode. Calibration of the standard agar pads was prepared as follows:

*Preparation of oxygen-free agar pad*: 1 g of sodium sulfite (Na_2_SO_3_) and 50 μL cobalt nitrate (Co(NO_3_)_2_) standard solution (ρ(Co) = 1000 mg L^-1^ in 0.5 M nitric acid) were dissolved in 100 mL lukewarm SM824 medium. Due to a spontaneous reaction of oxygen with the Na_2_SO_3_, the SM824 medium thereby becomes oxygen-free. Additional oxygen diffusing from air into the medium is removed by surplus Na_2_SO_3_. The oxygen-free medium was used to prepare 1-mm thick oxygen-free agarose pads (Ø = 18 mm, h = 1 mm) that were attached to the optodes for oxygen mapping. Images of the optode with the oxygen-free standard agar pads were taken by the oxygen detector unit (exposure 100,000 µm, gain 10).

*Preparation of an air-saturated agar pad*: 100 mL lukewarm SM824 medium was added to a 200-mL serum bottle placed in a 30^o^C water bath. To obtain air-saturated medium, air was blown into the stirred medium using an air pump with a glass-frit (air stone) to create fine air bubbles. After 20 min the pump was switched off and the medium was stirred for another 10 min at 30^o^C to avoid supersaturation of the medium. The medium was used to prepare air-saturated agarose pads (Ø = 18 mm), which were attached to the oxygen optode in the aforementioned microcosm. Images of the optode with the air-saturated standard agar pads were taken by an oxygen detector unit (exposure 100,000 µm, gain 10).

*Calibration curve*: A calibration curve (Figure S2) was created using signals measured in the oxygen-free and air-saturated agar pads following the instructions in the VisiSens AnalytiCal 1 software manual for calibration.

**Microbial cell recovery**

The bacterial cells in the agarose pad (cf. Fig. 1) were scraped with a blunt blade and transferred to 0.5 mL SM824 medium in 2-mL centrifugation tubes (Eppendorf, Hamburg, Germany). The agarose pad was further rinsed three times with 0.2 mL SM824 medium to collect remaining cells on the agar surface. The culture was then thoroughly vortexed for 40 s and used for total and viable cell counting.

**HPLC analysis**

Water-soluble fermentation metabolites (butyrate and 1-butanol) in the agarose pads (Fig. 1b) inoculated with *C. acetobutylicum* and *C. cinerea*, with *C. cinerea* only, with *C. acetobutylicum* only, or without inoculum were examined. Each agarose pad was divided into three parts with a spatula and weighed. Then, each pad fragment was mashed and mixed with 0.5 mL deionized water. Fermentation metabolites were extracted by intensive mixing of the agar-water samples for 15 s (Vortex Genie 2, Scientific Industries), followed by a 1 h period without mixing at room temperature (three repetitions). Afterward, the samples were centrifuged for 10 min at 20,800 × g and 4°C. The supernatants were filtered using syringe filter units with cellulose acetate membranes (0.2 μm in pore size; Labsolute) before measurement. The fermentation products were analyzed using a high-performance liquid chromatograph (Shimadzu Corporation) as described by Apelt (1) using a modified method with a column temperature of 55°C and a flow rate of 0.7 mL min^-1^.

**Calibration of oxygen beads**

Oxygen-free and air-saturated agarose pads were prepared as described above. The oxygen-sensitive beads were dispersed (CPOx beads, Colibri Photonics, Germany) in deionized water and 1 µL of the beads solution (~1200 beads µL^-1^) was placed to the oxygen-free or air-saturated agarose pad, respectively. After 5 min drying in a laminar flow, the agarose pads were flipped over and attached to the glass bottom of a Petri dish. Phosphorescence lifetimes of individual oxygen beads exposed to oxygen-free and air-saturated agarose pad were measured using the aforementioned OPAL-microscope system (cf. manuscript *Hyphal oxygen mapping by lifetime-based oxygen sensitive beads*). An average lifetime of 5.8 ± 0.2 s (*n* = 32 beads) and 4.6 ± 0.2 s (*n* = 30 beads) was measured in the air-saturated and oxygen-free agarose pad at 30^o^C, respectively. A conventional two-point calibration was done and calibration data was used for the hyphal liquid film oxygen measurement.

**Analysis of active swimming of *C. acetobutylicum* at different oxygen levels**

*100% air-saturated medium*: 80 mL SM824 medium were added to a 200-mL serum bottle. To obtain air-saturated medium, air was blown for 20 min into stirred medium using an air pump with a glass-frit (air stone), and the medium was stirred for another 10 min to avoid oversaturation of the medium.

*Oxygen-free medium*: 80 mL SM824 medium was added to a 200-mL serum bottle, and N_2_ was continuously flushed into the medium for 40 min to remove oxygen in the medium.

*80% air-saturated medium*: 1.6 mL of the 100% air-saturated medium and 0.4 mL of the oxygen-free medium were added to a 2-mL Petri-dish to obtain 80% air-saturated medium. The oxygen level in the medium increased from 80% to 82% air saturation after 20 min.

*Microscopic examination of* C. acetobutylicum *swimming activity*: 20 µL of the *C. acetobutylicum* culture with an OD_600_ = 1.1 was inoculated into the 2-mL Petri dish containing either air-saturated medium or 80% air-saturated medium. Swimming activity was examined microscopically at 5 min intervals for 30 min (Table S1). Experiments were performed in triplicates.

**Estimation of the effects of fungal activity on oxygen depletion**

Equations described by Oostra et al. (eqs. S1 and S2) (2) estimating oxygen penetration depth in solid-state fermentation (SSF) were used to estimate the maximal fungal biomass (*P_x_*) required to create anoxic conditions at 1-mm depth in liquid-filled upper soil layers.

$\delta= \left( \frac{2\cdot D_{e}{\cdot C}_{O,S}}{-r_{O}'''} \right)^{0.5}$	(eq. S1)

$-r_{O}'''= \frac{\mu\cdot P_{x}}{Y_{X/O}}$	(eq. S2)

In eq. S1, *δ* is the oxygen penetration depth (here *δ* = 1 mm), *D_e_* is the diffusion coefficient for oxygen in liquid-filled soil layer. *C_O,S_* is the saturated oxygen concentration in the liquid-filled soil layer at the soil-air interface. *-r_O_’’’* is the volumetric oxygen consumption rate and can be related to the fungal biomass using eq. S2, where *µ* is the specific fungal growth rate (h^-1^), *P_x_* is the mass fraction of the fungal biomass in the soil layers, *Y_X/O_* is the biomass/oxygen yield coefficient. To predict the maximal attainable oxygen flux in the liquid-filled soil layer, the oxygen diffusion coefficient in water at 30^o^C (*D_e_* = 3.7 × 10^‑9^ m^2^ s^‑1^) and saturated oxygen concentration in water at 30^o^C (*C_O,S_* = 0.22 mol m^-3^) were used. With the above parameters, to deplete the diffusing oxygen at *δ* = 1 mm depth in the water-filled soil layer, a maximal volumetric oxygen consumption rate *-r_O_’’’* = 1.628 × 10^‑3^ mol m^-3^ s^-1^ was required by calculation. Using a growth rate *µ* of 0.3 h^-1^ for soil fungi (3), a biomass/oxygen yield coefficient *Y_X/O_* = 1.16 Cmol mol^-1^ and a fungal biomass molecular weight of 0.024 kg C mol^-1^ (4), a fungal biomass of *P_x_* = 543 µg g^-1^ soil was calculated to deplete oxygen at 1-mm depth in liquid-filled soil layers.

**Table S1. Fraction of actively swimming *C. acetobutylicum* cells in liquid SM824 medium at different oxygen levels.** The fraction refers to the % of *n* = 249-281 cells detected in the focal plane during microscopic observation of triplicate samples.

| **Time of exposure**  (min) | **Percentage of actively swimming cells** | |
| --- | --- | --- |
|  | 80% air saturation | 100% air saturation |
| ~1 | 2.1 ± 0.2% | 0.0 ± 0.0% |
| 5 | 1.6 ± 0.2% | 0.0 ± 0.0% |
| 10 | 1.0 ± 0.2% | 0.0 ± 0.0% |
| 15 | 1.0 ± 0.2% | 0.0 ± 0.0% |
| 20 | 0.9 ± 0.2% | 0.0 ± 0.0% |
| 30 | 0.8 ± 0.0% | 0.0 ± 0.0% |

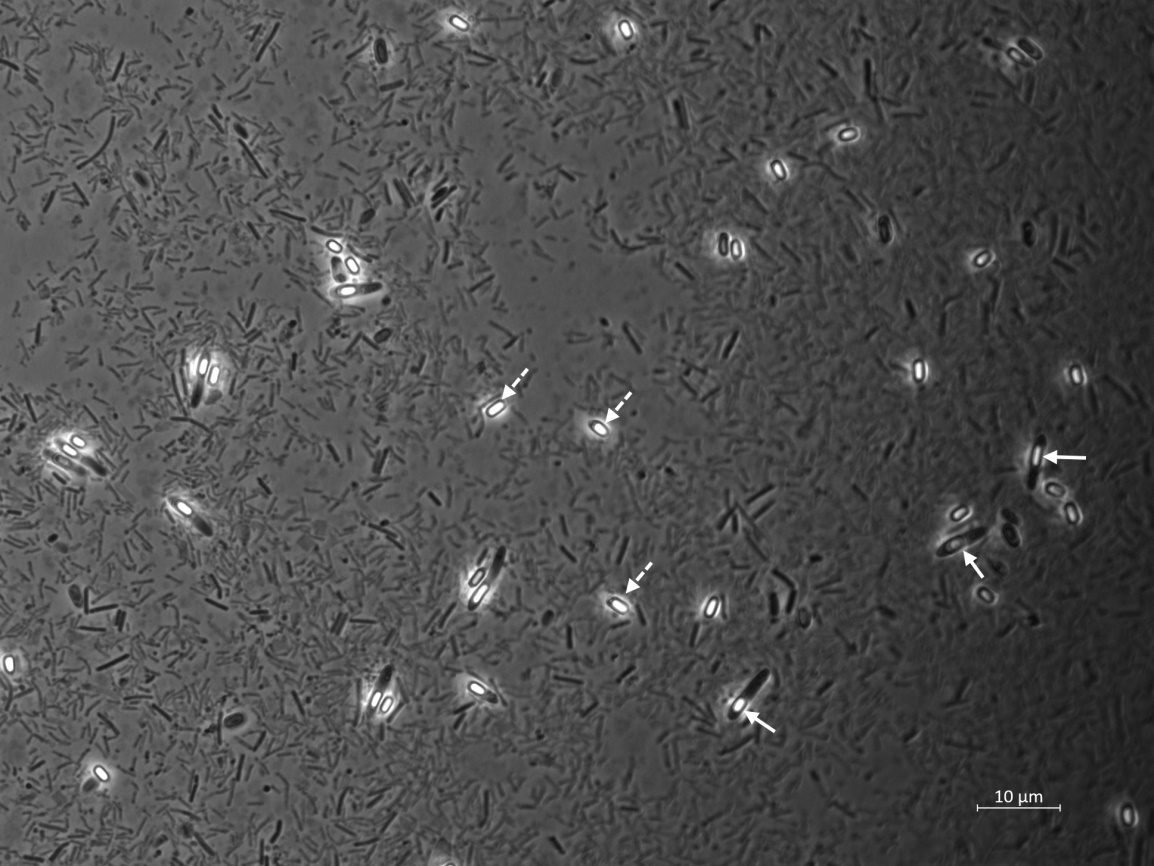

**Figure S1. Phase contrast micrograph of *C. acetobutylicum* spores**. White dashed arrows point at spores released from mother cells. White solid arrows highlight endospores in the swollen, cigar-like mother cells.

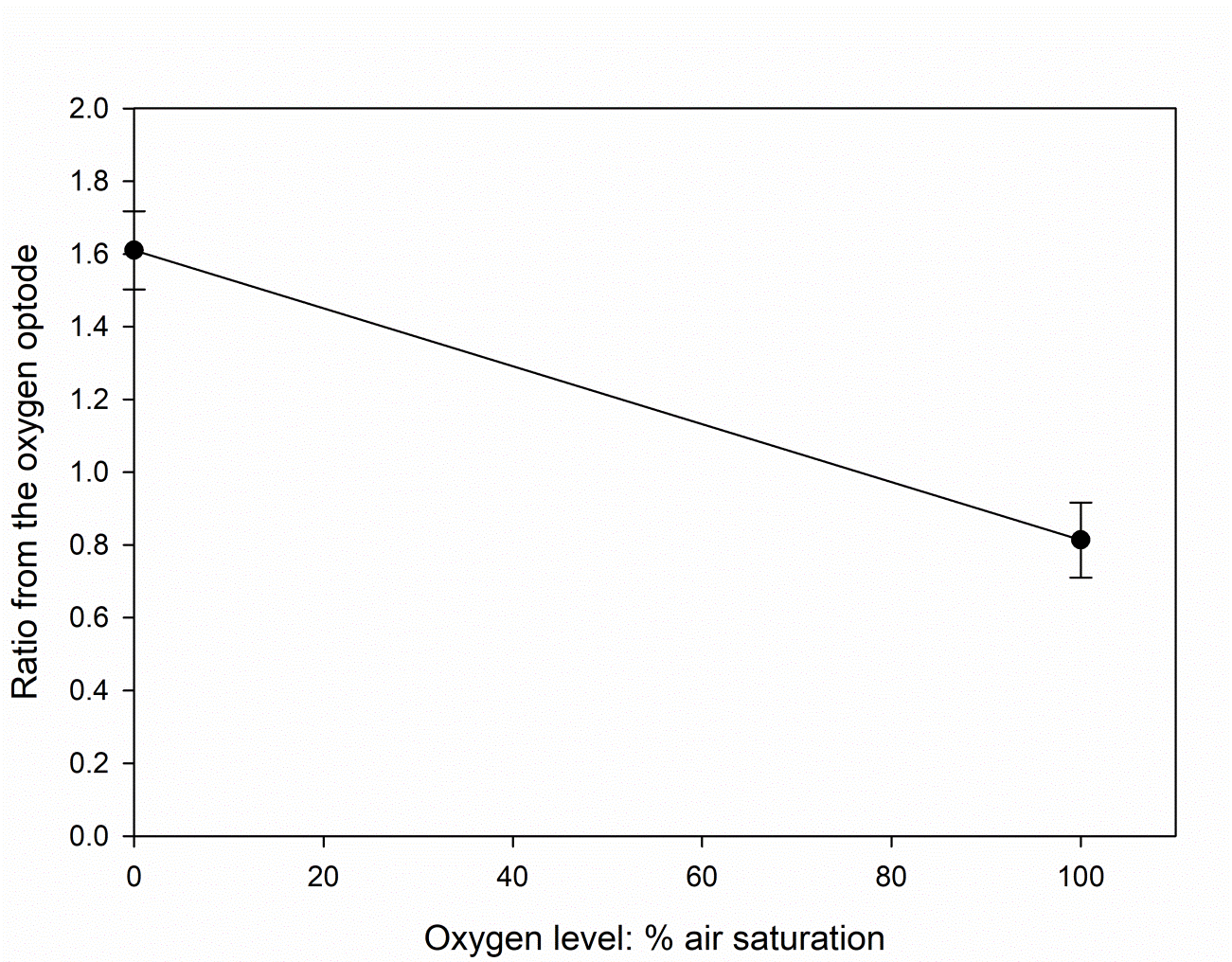

**Figure S2**. **Two-point calibration of the oxygen optode below an oxygen-free and an air-saturated agarose pad, resp**. Signals were measured in an area of 1.8 × 2.5 mm at each oxygen level at three spots. Ratiometric images were taken with a commercial detector unit using excitation and emission wavelengths that are not disclosed by the manufacturer.

.

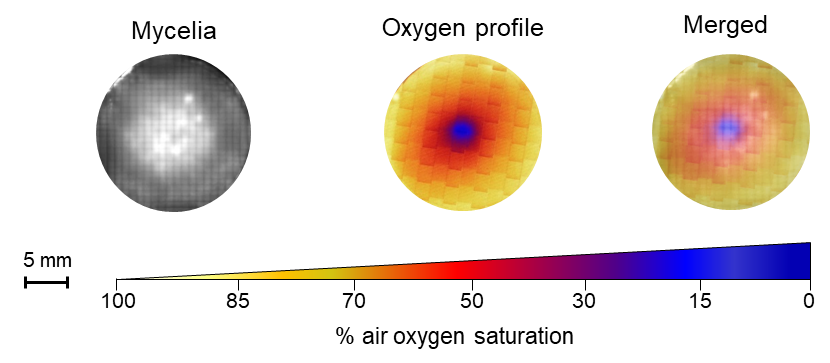

**Figure S3. Mycelial distribution and mycelial *in vivo* oxygen profiles of *C. cinerea* after 200 h of incubation**. The oxygen-depleted zone had a diameter of ca. 2.3 ± 0.6 mm.

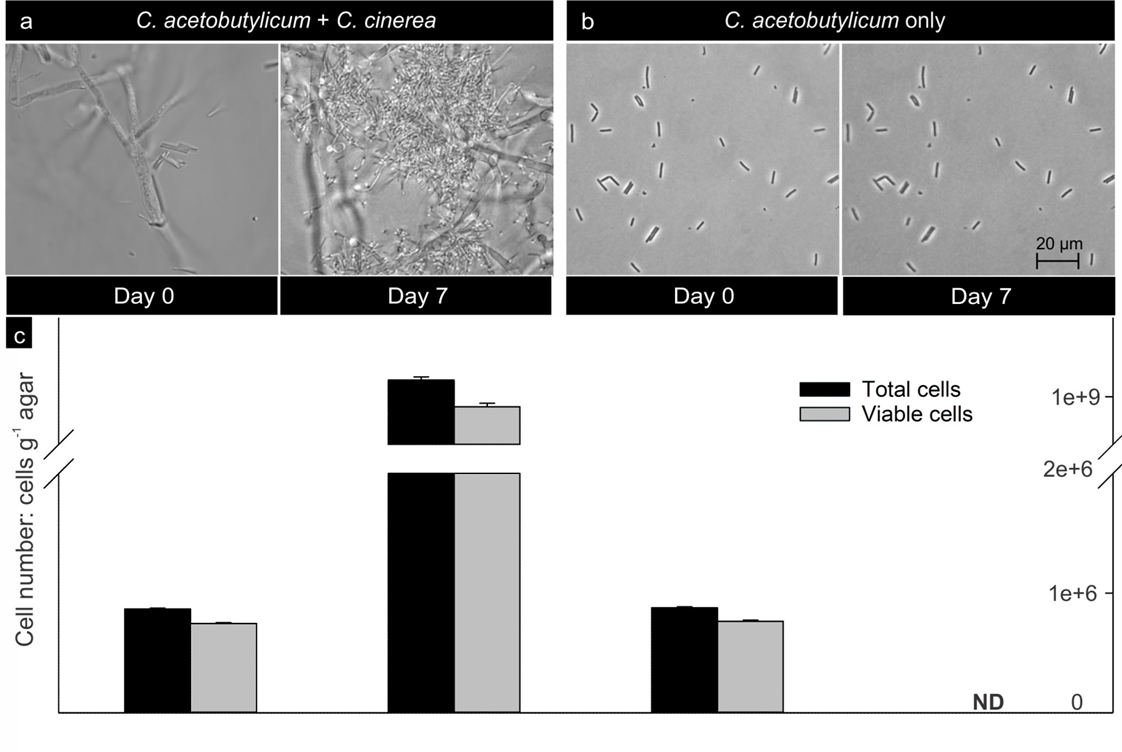

**Figure S4. *C. acetobutylicum* cell development after seven days in presence and absence of hyphae of *C. cinerea.* a & b** Micrograph of *C. acetobutylicum* cells after seven days in presence (**a**) and absence (**b**) of *C. cinerea.* **c)** *C. acetobutylicum* cell numbers density on the day of inoculation (t = 1 d) and on the day of harvest (t = 7 d) in the presence and absence of the fungus *C. cinerea*. Data represents the average and standard deviation (*n* = 3) of total and viable *C. acetobutylicum* cell counts. ND = not detected.

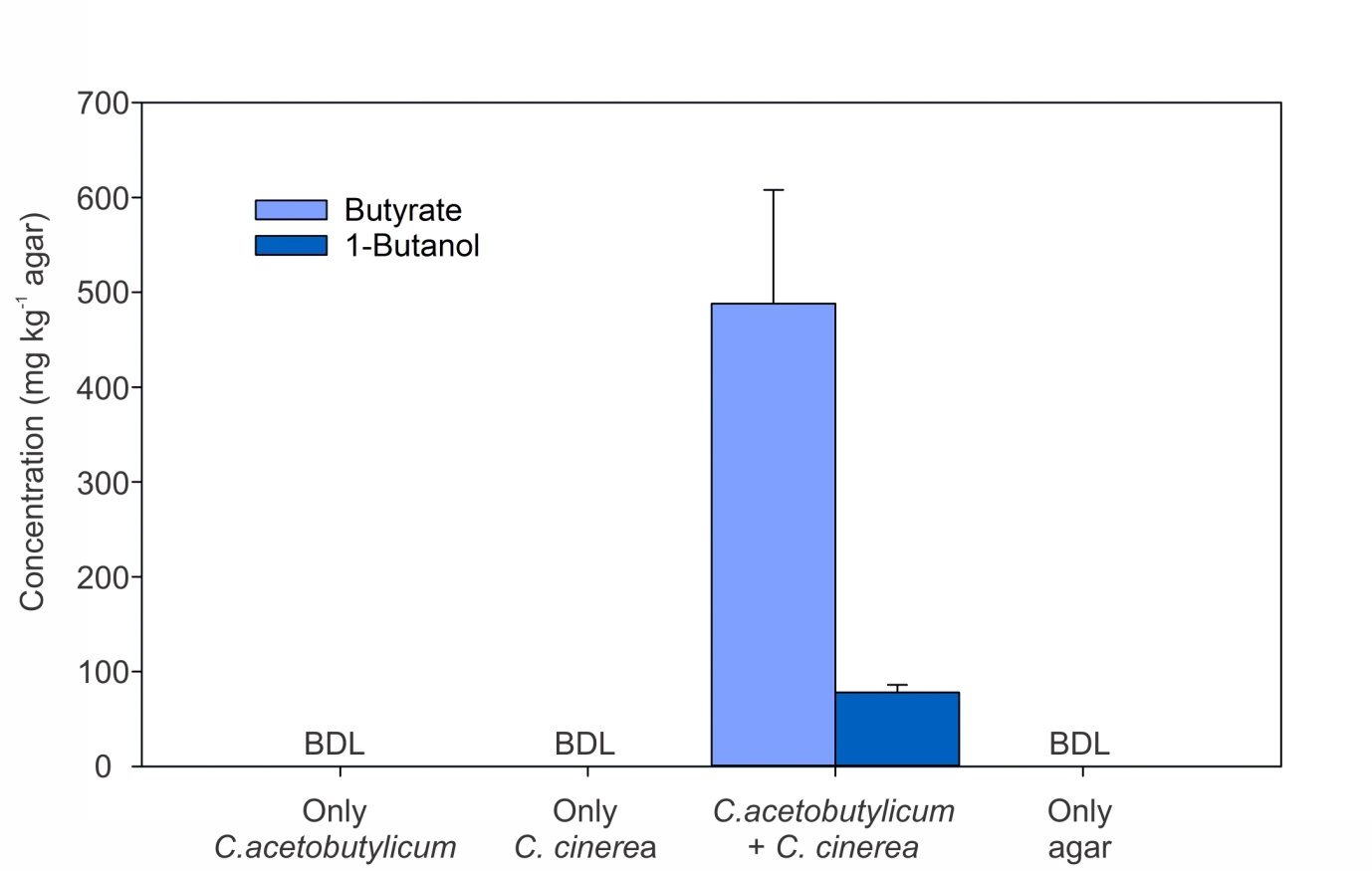

**Figure S5. Concentration of butyrate and 1-butanol in agarose pads with different inocula.** Butyrate and 1-butanol were solely detected in agarose pads inoculated with *C. acetobutylicum* and *C. cinerea* growing under ambient oxic conditions. Data represent mean and standard deviation of *n* = 2 for agar pads inoculated with *C. acetobutylicum and C. cinerea*, and *n* = 3 for the treatments with *C. acetobutylicum* only, *C. cinerea* only or agar only. BDL = below detection limit.

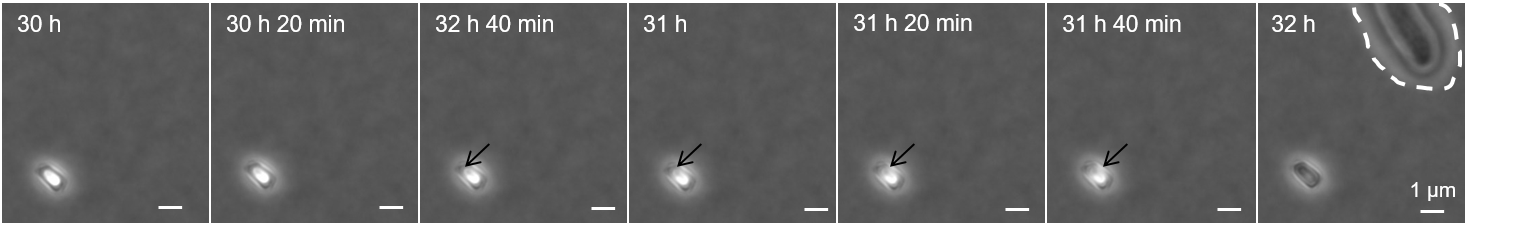

**Figure S6**. **Spore development of *C. acetobutylicum* two hours before appearance of hyphae of *C. cinerea*.**

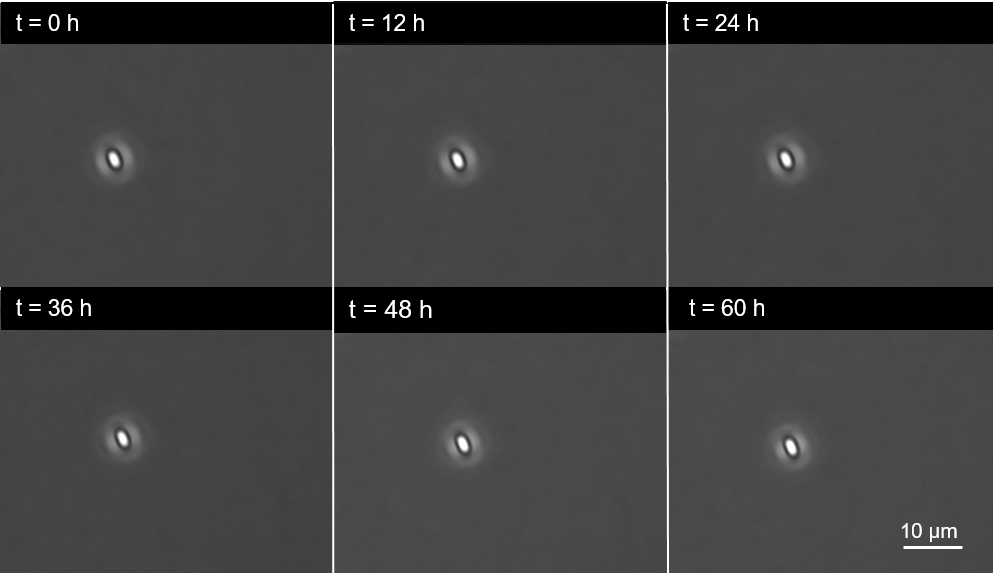

**Figure S7.**  ***C. acetobutylicum* spores in the absence of *C. cinerea* mycelia exposed to oxic condition over 60 h of incubation.**

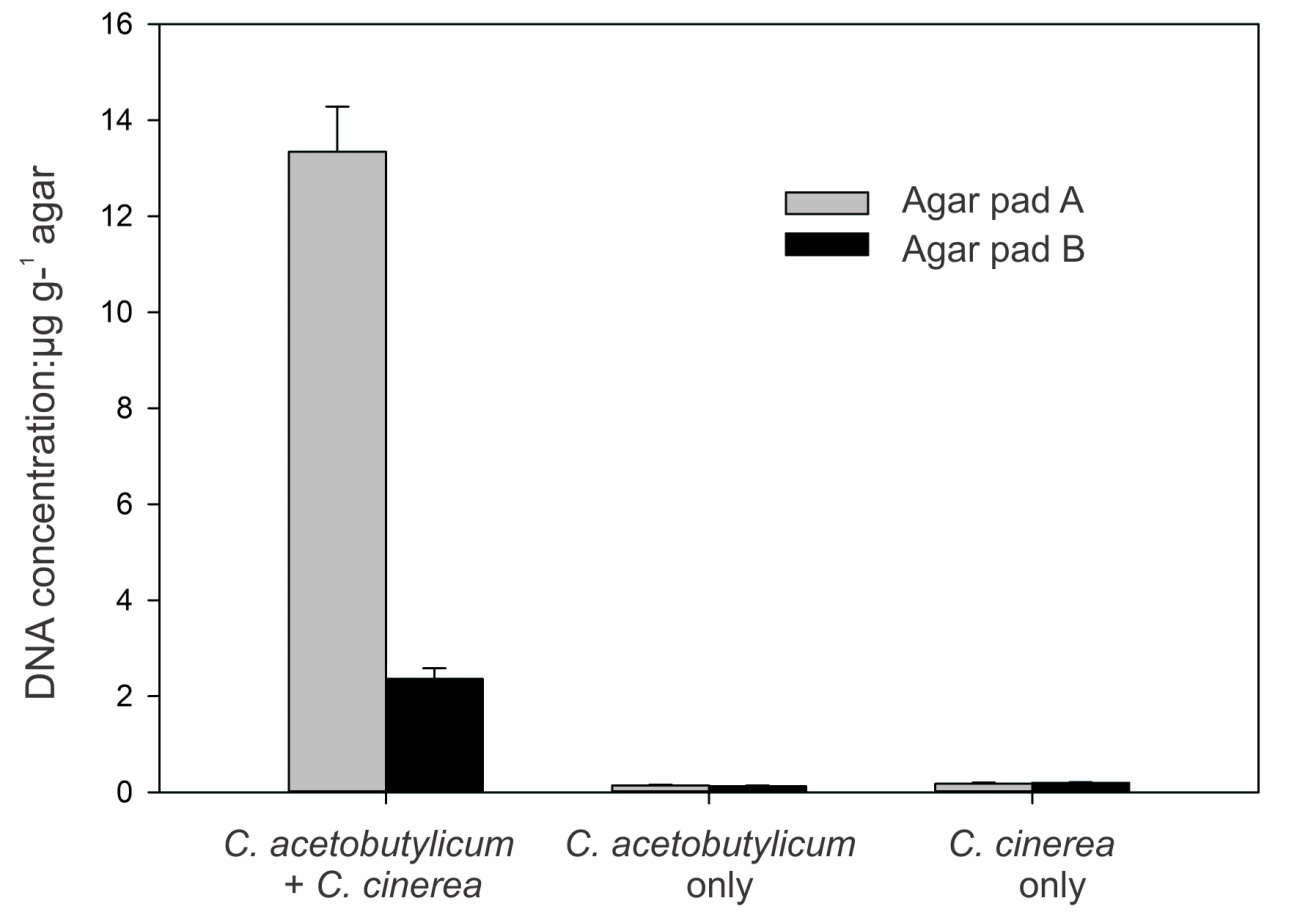

**Figure S8. Extracted DNA after seven days in agar pads A and B in presence of (i) *C. acetobutylicum* and *C. cinerea*, (ii) *C. acetobutylicum* and (iii) *C. cinerea.*** Data represent the average and standard deviation of *n* = 3 DNA extractions.
